# Characterizing the monomer-dimer equilibrium of UbcH8/Ube2L6: A combined SAXS and NMR study

**DOI:** 10.1101/2023.04.13.536743

**Authors:** Kerem Kahraman, Scott A. Robson, Oktay Göcenler, Cansu M. Yenici, Cansu D. Tozkoparan, Jennifer M. Klein, Volker Dötsch, Emine Sonay Elgin, Arthur L. Haas, Joshua J. Ziarek, Çağdaş Dağ

**Affiliations:** Nanofabrication and Nanocharacterization Center for Scientific and Technological Advanced Research (n2STAR), Koc University, İstanbul, Turkey; Department of Pharmacology, Feinberg School of Medicine, Northwestern University, 320 East Superior Avenue, Chicago, IL, 460611, USA; Department of Biochemistry and Molecular Biology, LSUHSC-School of Medicine, 1901 Perdido Street, New Orleans, LA, 70112, USA; Centre for Biomolecular Magnetic Resonance, Institute for Biophysical Chemistry, Goethe-University of Frankfurt/Main, Germany; Muğla Sıtkı Koçman University, College of Sciences, Department of Chemistry, Muğla, 48000, Turkey; Koc University Isbank Center for Infectious Diseases (KUISCID), Koc University, Istanbul, Turkey

**Keywords:** . ISGylation, ubiquitination, E2 enzyme, TRACT, oligomerization

## Abstract

Interferon-stimulated gene-15 (ISG15) is an interferon-induced protein with two ubiquitin-like (Ubl) domains linked by a short peptide chain, and the conjugated protein of the ISGylation system. Similar to ubiquitin and other Ubls, ISG15 is ligated to its target proteins through a series of E1, E2, and E3 enzymes known as Uba7, Ube2L6/UbcH8, and HERC5, respectively. Ube2L6/UbcH8 plays a literal central role in ISGylation, underscoring it as an important drug target for boosting innate antiviral immunity. Depending on the type of conjugated protein and the ultimate target protein, E2 enzymes have been shown to function as monomers, dimers, or both. UbcH8 has been crystalized in both monomeric and dimeric forms, but the functional state is unclear. Here, we used a combined approach of small-angle X-ray scattering (SAXS) and nuclear magnetic resonance (NMR) spectroscopy to characterize UbcH8’s oligomeric state in solution. SAXS revealed a dimeric UbcH8 structure that could be dissociated when fused N-terminally to glutathione S-transferase. NMR spectroscopy validated the presence of a concentration-dependent monomer-dimer equilibrium and suggested a backside dimerization interface. Chemical shift perturbation and peak intensity analysis further suggest dimer-induced conformational dynamics at E1 and E3 interfaces - providing hypotheses for the protein’s functional mechanisms. Our study highlights the power of combining NMR and SAXS techniques in providing structural information about proteins in solution.

## INTRODUCTION

Interferon-Stimulated Gene 15 (ISG15), also known as hUCRP or IP17, is a 15 kDa ubiquitin-like, type I interferon (IFN) inducible protein [1]. ISGylation is a ubiquitin-like (Ubl) post-translational modification (PTM) that involves the covalent attachment of ISG15 to target proteins [2]. Similar to other Ubls, ISGylation plays important roles in various cellular processes such as innate antiviral immunity, protein degradation, and signal transduction [3]. Free, unconjugated, ISG15 also serves immunoregulatory functions as a cytoplasmic and secreted signaling protein in eukaryotic organisms [4]. Inherited ISG15 deficiency dramatically reduces the innate immune system’s ability to fight viruses in mice yet only appears to cause immunoregulatory issues against mycobacterial, not viral diseases, in humans [3]. Thus, the role of ISG15 in human viral pathogenesis is not clearly understood.

The ISGylation cascade requires the sequential action of three enzymes: Ube1L as the E1 enzyme, UbcH8 as the E2 enzyme, and HERC5 as the E3 enzyme. First, ISG15 binds the catalytically active cysteine of the Ube1L activating enzyme (E1) in an ATP-dependent reaction. Then, E1 interacts with UbcH8 conjugating enzyme (E2) through its ubiquitin folding domain (UFD), which facilitates the transesterification of active ISG15, and results in an intermediate ISG15-UbcH8 complex joined by a thioester bond [5]. Finally, HERC5 ligase enzyme (E3) interacts with the intermediate ISG15-UbcH8 complex to mediate ligation of ISG15 to the target protein. UbcH8 plays a central role in ISGylation as it interacts with both E1 and E3 enzymes - making it a key target for the regulation of the ISGylation pathway [6].

Under reducing conditions, E2 enzymes can spontaneously form dimers when a crosslinker is added [7], and apart from a few exceptions, E2 enzymes are capable of preserving their dimer form [8, 9]. Both the dimer and the monomer forms of E2 enzymes are capable of recruiting E3 enzymes and conjugating ubiquitin [10]. Although dimeric E2 enzymes are perceived as more advantageous because one of the monomers can remain associated while the ubiquitin conjugation continues with the other.

The Protein DataBank (PDB) contains both dimeric (PDB ID:1WZV) and monomeric (PDB ID:1WZW) crystal structures of UbcH8 (Supplementary Figure 1). Yet, it is unknown whether UbcH8 dimerizes naturally or as a consequence of non-specific crystal packing contacts. In this study, we aimed to characterize the oligomeric state of human UbcH8 (Ube2L6) in solution using Small Angle X-ray Scattering (SAXS). We first used a fusion protein approach with the goal of producing a high-resolution scattering envelope to properly place the UbcH8 protomers. Surprisingly, upon removal of the N-terminal fusion, monomer-dimer equilibrium further shifted to the dimer side. We next used solution nuclear magnetic resonance (NMR) spectroscopy to first validate and then further characterize the monomer-dimer equilibrium. Our results indicate that UbcH8 contains a substantial dimer population at 300 μM concentration and that dimerization may induce conformational changes at the distal E1 and ISG15/E3 interaction interfaces.

## RESULTS

### GST fusion guides SAXS protein structural modeling

To determine the state of UbcH8 in solution, we expressed and purified it fused to an N-terminal glutathione-S transferase (GST) tag herein termed GST-UbcH8. We hypothesized that the 28 kDa GST molecule should be easily discernible from the smaller (18 kDa) UbcH8, and would dramatically improve fitting SAXS scattering data to the structural model. The sample was concentrated to 300 μM and six, 10 min SAXS frames were collected for a total of one hour. Superimposition of all 10 min frames confirmed that the X-ray beam produced little to no detectable radiation damage (Supplementary Figure 2). The medium to high q region, which is emphasized in the q vs l(q) plot, is consistent with a folded sample (Figure 1A). The Kratky plot possessed a bell-shaped curve that approaches zero after reaching a maximum at ∼3 sRg; this result is consistent with a properly folded, globular protein (Figure 1B). While slight deviations between the typical Kratky plot and dimensionless Kratky plot can aid in the assessments of flexibility, no apparent differences were observed.

**Figure 1.**
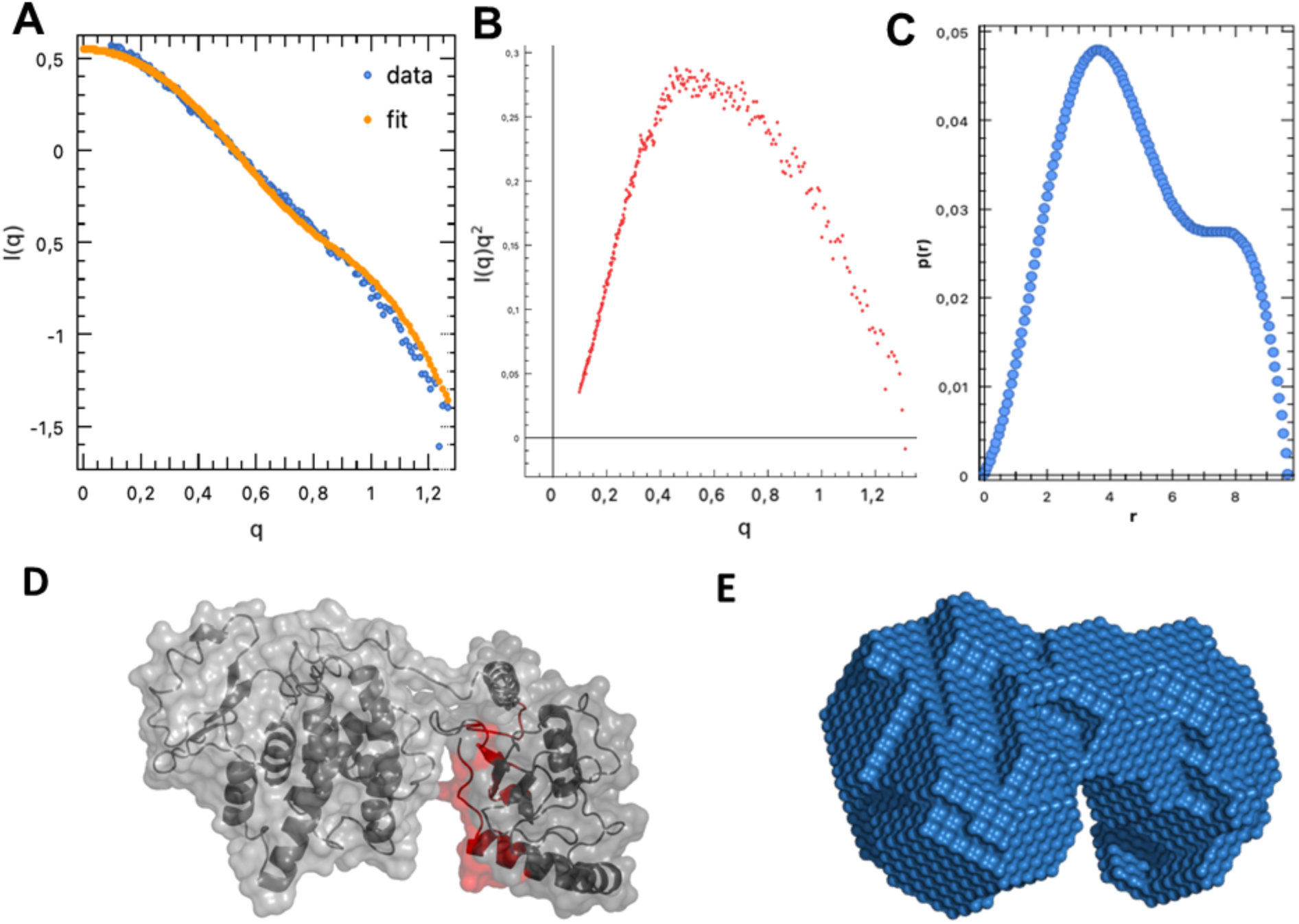
SAXS analysis of GST-UbcH8 fusion protein. (A) ln I (s) vs s plot, (B) Kratky plot, and (C) Pair distance distribution, P(r), plot of the experimental SAXS intensity obtained at 4.1 mg/ml (300 μM) of the GST-UbcH8 fusion protein. The pair-distance distance distribution plot of the GST-UbcH8 fusion protein scattering data calculated by GNOM. (D) The individual crystal structures of GST (left) (PDB:1R5A) and UbcH8 (right (PDB:1WZW) shown as cartoon representations; UbcH8 homodimeric dimerization surface labeled as red. (E) The GST-UbcH8 dummy atom model, obtained by the ATSAS online package, is shown as a surface representation.

The pair-distance distribution function, P(r), is a measure of the frequency of interatomic distances that can also provide information about the protein shape. The presence of a shoulder in the P(r) suggests a multidomain protein as expected for the GST-UbcH8 fusion (Figure 1C). The largest distance (D_max_) in the P(r) histogram was 8 nm (Figure 1C). The GST-UbcH8 crystal structures were then fitted into the final 3D DAMMIF dummy atom model (Figure 1D,E). In addition, Molecular Size Parameters obtained with Primus are included in Supplementary Table 1 and match the theoretically expected dimensions.

Both GST and UbcH8 proteins, as well as the linker peptide, are clearly visible fitting a monomeric model. The fact that even the linker region can be detected with SAXS analysis and be observed this clearly, underscores the power of SAXS in structure determination. After GST cleavage, we recorded unexpected UbcH8 size exclusion chromatographs. The UbcH8 peak was dispersed over a much wider elution volume compared to a similarly-sized (12.2 kDa) monomeric protein when injected using identical run parameters (Supplementary Figure 3) leading us to hypothesize that GST may block the dimerization site. Figure 1D is the GST-UbcH8 fusion surface representation using SWISS-MODEL; the UbcH8 homodimerization surface is shown in red. Both the SWISS-MODEL and the SAXS model positioned the GST fusion protein to occlude the UbcH8 dimerization interface.

### FreeUbcH8 is a Dimer in Solution

To test how well GST-fusion improves the modeling of UbcH8 into the SAXS scattering, we prepared a second UbcH8 sample with the GST protein removed. Again, we concentrated UbcH8 to 300 μM and collected six, 10 min frames for a total acquisition time of 1 h (Figure 2A). Similar to GST-UbcH8, the Kratky plot possessed a bell-shaped curve that approaches zero (Figure 2B). We estimated a slightly larger R_g_ ∼ 4.40 nm, compared to GST-UbcH8, from the low q region, whereas the P(r) D_max_ was reduced to 6.2 nm (Figure 2C). Surprisingly, the free UbcH8 P(r) also contained a shoulder suggesting homodimerization (Figure 2C). ATSAS molecular weight analysis predicts a 39.5 kDa particle, which is approximately double the expected 18 kDa UbcH8. We then fitted the UbcH8 dimer crystal structure (PDB 1WZV; Figure 2D) to the dummy atom model of the scattering envelope (Figure 2E). The best-fit model (X^2^ = 1.7) possesses a dimerization interface with the active site cysteines (Cys85) of each protomer pointed outwards (Figure 2D,E). The consistency between the previously published dimer crystal structure and the dummy atom model obtained by SAXS analysis supports UbcH8 homodimerization in the absence of a GST-tag.

**Figure 2.**
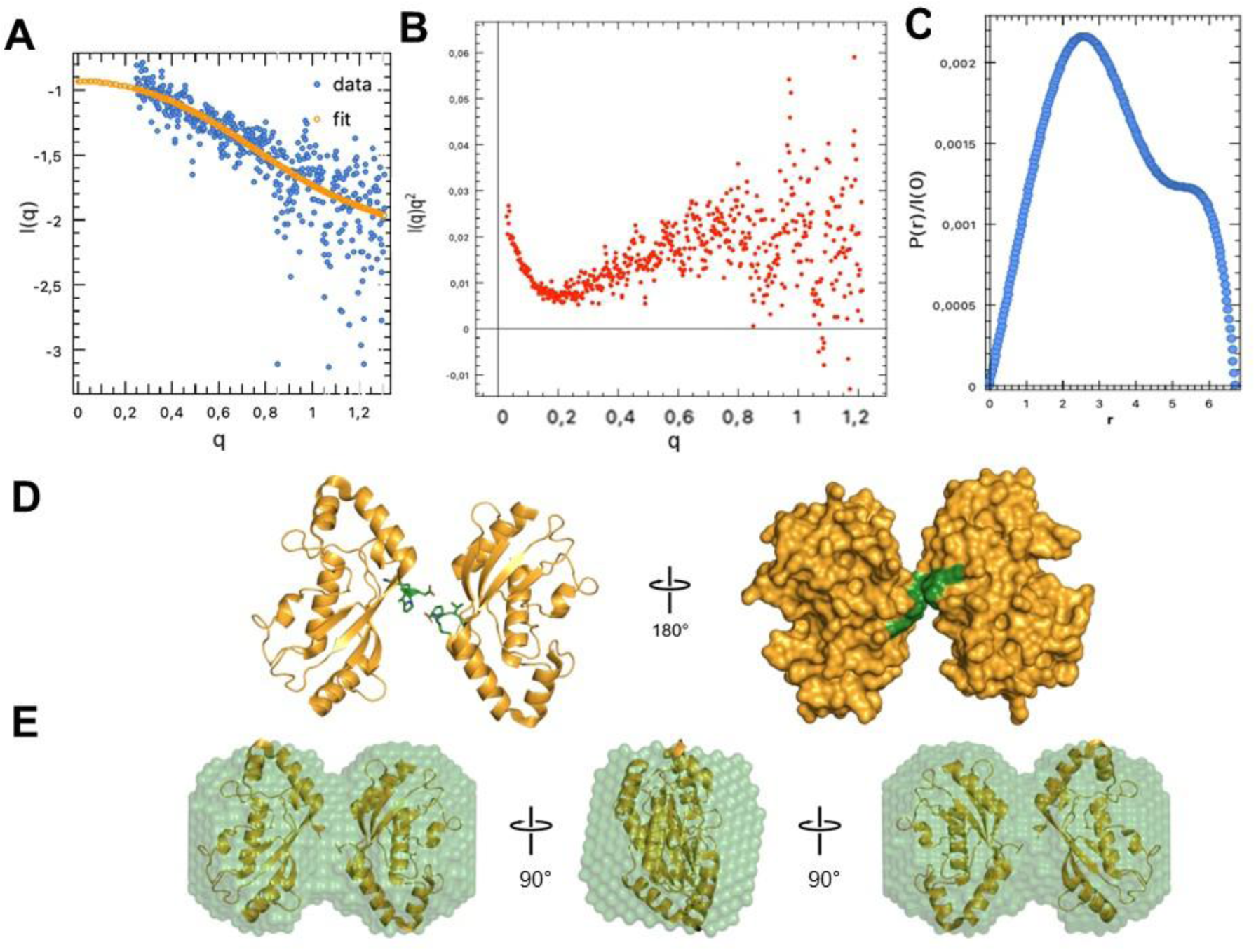
SAXS analysis of free UbcH8. (A) ln I (s) vs s plot, (B) Kratky plot, and (C) Pair distance distribution, P(r), plot of the experimental SAXS intensity obtained at 4.1 mg/ml (300 μM) free UbcH8 protein. (D) Crystal structure of UbcH8 dimer (pdb:1WZV) shown as cartoon and surface representation using Pymol. Residues D149, R150, and P151 of the dimerization interface are colored forest green. (E) UbcH8 dimer crystal structure (pdb:1WZV) fitted (χ = 1.02) into the DAMMIF dummy atom model using SASpy.

### NMR analysis of UbcH8 monomer-dimer equilibrium

To further establish dimerization of UbcH8 in solution, we performed Transverse Relaxation Optimized Spectroscopy (TROSY) for rotational correlation times (TRACT) experiments [11,12] to estimate the rotational correlation time (*π_c_*) of UbcH8 at two different concentrations: 300 μM and 150 μM (Figure 3). The signal intensity ranging from 8.6 to 9.2 ppm was integrated to maximize signal to noise and emphasize well-structured regions of the protein that are representative of global tumbling. We estimated ^15^N relaxation rates for the TROSY and anti-TROSY integrated signals using Bayesian Parameter Estimation of a two-parameter single-exponential decay model. This method produces a distribution of decay rates, which encompass uncertainty, that were then used to determine the cross-correlated relaxation (CCR) rate. The rotational correlation time was estimated from CCR according to an algebraic solution [12] of the modified Goldman relation [13], assuming an order parameter (*O^2^*) of 0.8. We determined a *π_c_*∼16 ns at 300 μM and ∼13 ns at 150 μM (Figure 3), which demonstrates a concentration dependence on molecular rotation diffusion times. We then used hydroNMR [14] to model rotational diffusion of monomeric and dimeric UbcH8 from the PDB 1WZV dimeric crystal structure. hydroNMR reported a *π_c_*= 20.5 ns for the dimer and 7.4 ns for the monomer at 25 °C. Taken together, this confirms that UbcH8 undergoes monomer-dimer exchange and indicates a substantial dimer population even at 150 μM. Data could not be collected at lower concentrations due to the sensitivity limit of the room temperature NMR probe.

**Figure 3.**
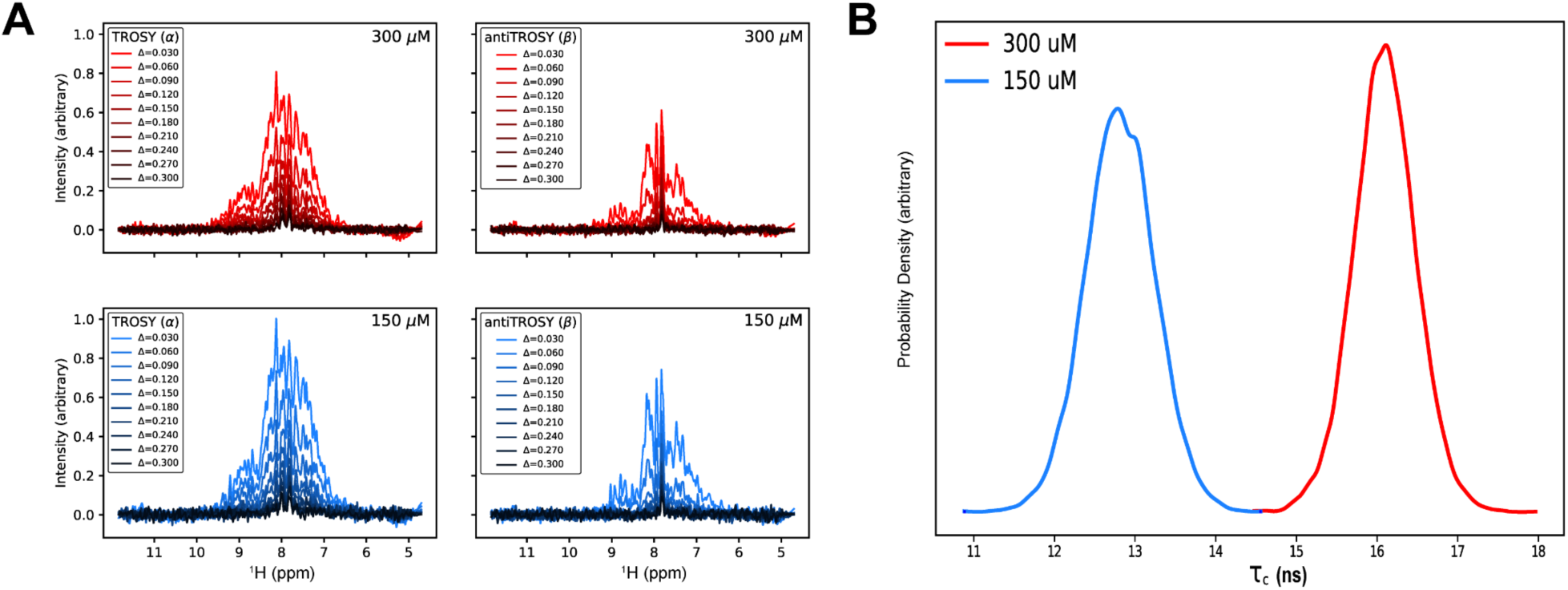
TRACT analysis of free UbcH8 to determine rotational correlation time. (A)The UbcH8 1D ^15^N TROSY (left) and anti-TROSY (right) spectra from the TRACT experiment. The top and bottom panels are UbcH8 at 300 (red) and 150 μM (blue), respectively. We integrate from 8.6-9.2 ppm (gray boxed region) under the assumption that it possesses resonances from primarily structured regions. (B) The probability density estimates of the overall rotational correlation time (*π_c_*) for UbcH8 at 300 μM (red) and 150 μM (blue). The average (point) estimate for 300 μM and 150 μM UbcH8 are 16.1 ns and 12.8 ns, respectively.

We next collected ^15^N heteronuclear single quantum coherence (HSQC) solution NMR spectra at 150 μM (Supplementary Figure 4) and 300 μM to identify UbcH8’s dimerization interface. Resonances were assigned by visual inspection using BMRB Entry ID 16321 as a reference list. The NH resonances of all residues except for the 18 prolines were assigned (86.18% completion). We then assessed both concentration-dependent chemical shift perturbations (CSPs) and peak intensity differences. Substantial concentration-dependent CSPs were induced at locations far from the crystallographic dimerization interface (Figure 4). All of the perturbed residues except for K17 and Y21 are situated at either the E1 or ISG15/E3 interaction surfaces. Residues S91, E92, and S97 are clustered on a loop near the catalytic C85 residue where ISG15 is covalently attached. Whereas S97, T100, V103, V111 and N115 are proximal to the ISG15/E3 binding region on the UbcH8 surface; interestingly, these residues are arranged towards the UbcH8 core rather than at the surface (Figure 4). Given that E1 and ISG15/E3 involve distinct interfaces, we hypothesize a conformational change or allosteric pathway influences the transfer or binding of ISG15. Our results suggest that dimerization may play an additional role in ISGylation. We hypothesize that the weak CSPs could reflect a mostly sidechain-mediated interface and/or that the ensemble is predominantly dimeric even at 150 μM concentration.

**Figure 4.**
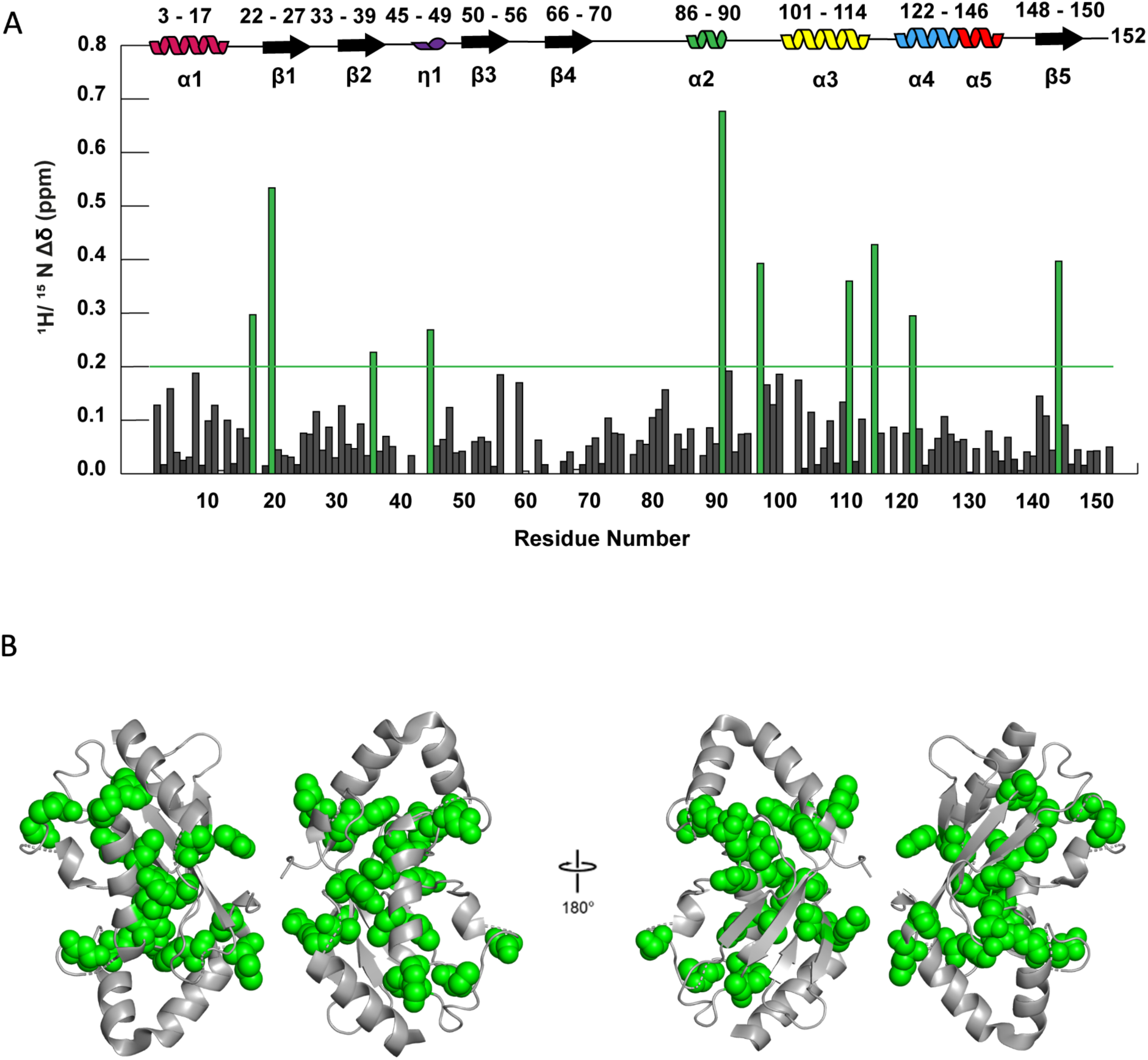
^1^H-^15^N Chemical shift perturbations mapped onto UbcH8 dimeric crystal structure. A) The combined ^1^H-^15^N chemical shift perturbations between 300 μM and 150 μM UbcH8 were calculated for each residue. The residues colored green possessed chemical shift perturbations larger than the threshold (red line). Residues with no bars were unobservable at either concentration. B) Residues with chemical shift perturbations larger than the threshold were mapped onto the UbcH8 dimeric crystal structure (PDB 1WZV). These residues cluster to two distinct regions: the E1 binding surface (K17, Y21, A36, Y45) and ISG15/E3 binding surface (S97, T100, V103, V111 and N115).

Thus, we also measured the concentration-dependent changes in peak intensity. We hypothesize these intensity differences result from monomer-dimer exchange on the intermediate (microsecond-millisecond) timescale, but it’s also possible that they reflect dimerization-dependent fluctuations in longitudinal (T_1_) or transverse (T_2_) relaxation. The largest changes in peak intensity again clustered to the E1 and ISG15/E3 interfaces, while also highlighting an extended region along the crystalized dimer interface (Figure 5). D149 sits at the center of the dimerization surface with E141 and L144 in close proximity. It’s plausible that D28 and A29, located in a loop region of the opposing protomer, could possess the flexibility to interact. Furthermore as the overall structure gets bigger with the dimerization, decreased signals from some peaks were expected due to line broadening. Although unlike the CSP analysis which showed that most of the conformational changes occurred away from the dimerization site, delta chemical shift intensity analysis revealed that most of the affected residues were on the dimerization site. In fact, the peak intensity of N23 and D149 residues from opposing protomers, which are within 3.3 Å distance in the crystal structure, deviated from the mean peak height by 44.7% and 62.2%, at 150µM, and 37.6% and 35.6%, at 300 µM, respectively (Supplementary Figure 5). Taken together, this indicates that dimerization is *ipso facto* involved in defining interaction dynamics between E1 and E2 enzymes.

**Figure 5.**
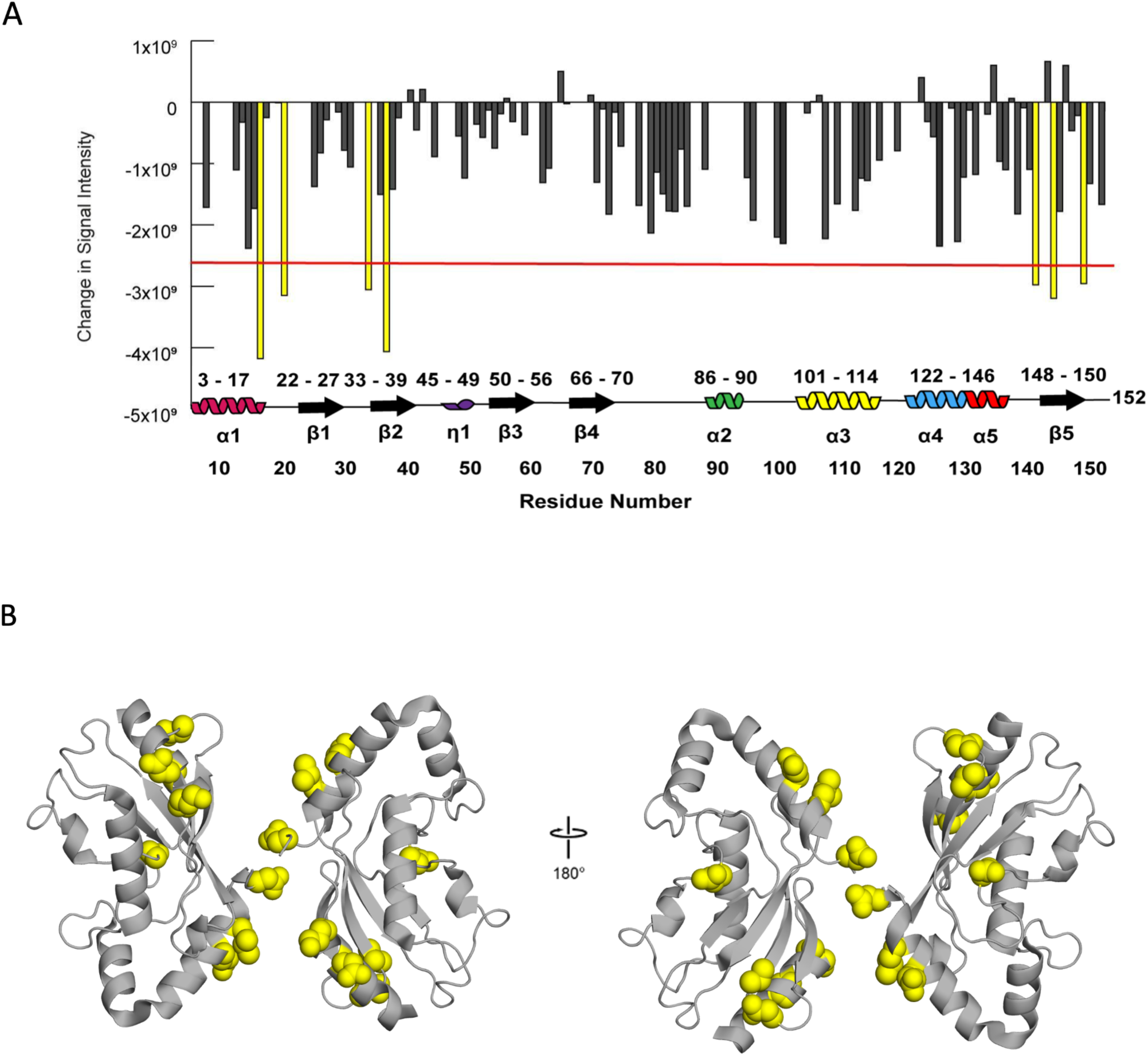
Concentration dependent chemical shift signal intensity changes. A) Concentration-dependent changes in ^1^H-^15^N peak intensity mapped onto UbcH8 dimeric crystal structure. The residues colored yellow possessed peak intensity changes that were larger than the threshold (red line); these residues are located at the E1 binding surface (D12, K16, N30, V33) and C-terminal (E141, L144, and D149). Residues with no bars were not observed at either concentration. B) Residues with peak intensity changes larger than the threshold were mapped onto the UbcH8 dimeric crystal structure (PDB 1WZV)

As a further attempt to investigate UbcH8 dimerization effect on non-covalent interactions between ISG15 and UbcH8, 260 uM ^15^N labeled UbcH8 was titrated with unlabeled ISG15. The chemical shift perturbations were mapped on the UbcH8 structure and were observed to affect different residues (Supplementary Figure 6) compared to the 150µM-300 µM UbcH8 spectra. The catalytic C85 residue to which ISG15 binds is located on the opposite side of the proposed dimerization site on the UbcH8. This binding surface is similar to the Ubiquitin binding surface on UbcH8 [15]. In addition, since we only examined non-covalent interactions, the changes in our CSP values are lower than the values obtained by covalently binding Ubiquitin to UbcH8 with a di-sulfide bond [15]. Based on the data, rather than dimerization inhibits ISG15 interaction, it is more likely that dimerization may affect E1 and/or E3 interactions, thus negatively regulating the ISG15 conjugation pathway.

## DISCUSSION

In this study, we investigated the oligomeric state of UbcH8 in solution using Small Angle X-ray Scattering (SAXS) and NMR analysis. To improve the fitting of SAXS scattering data to the structural model, we initially employed an N-terminal GST-fused UbcH8 protein. The results from SAXS experiments and analysis indicate that the GST-UbcH8 fusion protein is monodisperse and properly folded in solution. The 3D DAMMIF dummy atom model also revealed a monomeric model of the GST-UbcH8 fusion protein, highlighting the advantages of SAXS in structure determination. The free form of UbcH8, without the GST fusion, was further investigated. Surprisingly, the P(r) distribution suggested a multidomain complex, and the ATSAS molecular weight analysis indicated dimer formation. The consistency between the existing crystal structure of UbcH8 dimer and the SAXS-derived dummy atom model supports the formation of a UbcH8 homodimer in solution in the absence of a GST-tag.

Our NMR analysis, including chemical shift perturbation and peak intensity measurements, provide additional evidence for the dimerization of UbcH8. Residues involved in the dimerization process were identified, and the effects of dimerization on the E1 and E3 interaction sites were observed. The TRACT experiments also supported the dimerization of UbcH8, revealing a concentration-dependent behavior of UbcH8 in solution, which suggests monomer-dimer exchange on an intermediate timescale. The dimerization of UbcH8 and its implications on the ISGylation process are consistent with previous reports of E2 enzymes forming dimers to facilitate polyubiquitination [10]. Dimerization of several E2 enzymes are reported and the observed dimerization of the enzymes are found to be stimulating the catalytic activity of the E2 enzyme in these studies [9, 16, 17, 18]. In 2010, David et. al. suggested that E2 enzymes form dimers in solution regardless if an active ubiquitin is present [7]. While the monomer form is also active for the acquisition of ubiquitin, the dimer form of the E2 is found to be more advantageous as while one monomer site is binding the ubiquitin molecules, the other site is capable of remaining associated to the target protein and thus facilitates efficient polyubiquitination. The acting mechanism of E2 enzymes proposed in this study suggests that the E2 enzymes function as dimers while catalyzing polyubiquitination process [7]. An example of dimerization is observed for the E2 enzyme UBE2W [8]. An equilibrium between the monomeric and dimeric states was observed and verified using biochemical assays as well as NMR experiments. To compare the type of UbcH8 dimerization with previous dimer E2 structures, we screened the fitting of free UbcH8 dummy atom model with previously-determined dimeric E2 enzyme crystal structures (Supplementary Figure 7 and Supplementary Table 2). These structural screening results showed us that the dimerization pattern of UbcH8 is most similar to UbcA1 from Ube2w family (Supplementary Figure 8).

In another study conducted on the Ub E2 enzyme UBE2S, intracellular colocalization was observed and the presence of oligomeric states was measured using NMR [9]. Additionally, the authors propose the dimerization model where the C-terminal helices of the protein monomers “hug” the other, thus generating a dimer that is not auto ubiquitinated as opposed to the monomeric form. Prevention of auto ubiquitination allows UBE2S to be present at higher intracellular concentrations without being degraded by the proteasome, although its ubiquitination activity is impaired. This mechanism shows how dimerization could be relevant for regulation of ubiquitination activity. As reviewed by [6], the ubiquitination activity of certain E2s can also be modulated by the binding of Ub to a secondary site, named the backside, different from the catalytic surface. For different E2 enzymes, the poly Ub chain formation capability can be enhanced or impaired by the binding of Ub to the backside surface. Additionally, the reported binding of E3 enzymes to the backside [6] also raises the question of how ubiquitination activity is regulated by the equilibrium between E1-E2 and E2-E3 interactions and how E2 dimerization could affect this process. The complex interaction dynamics of E2 with itself, the conjugated protein, and the E3 enzymes are yet to be revealed as well as how dimerization affects this regulation. Dimerization could reduce E2 activity as in the case of UBE2W, or by limiting backside accessibility to E3 or the conjugate Ubl protein. Contrarily, E1 and E3 interaction surfaces on the backside could be exposed more, leading to increased E2 activity. Overall, E2 dimerization holds significance for regulation that is yet to be revealed.

ISG15 and related enzymes are expressed at elevated levels when an infection is present, or when stimulated by IFNs. SARS-CoV-2 infected macrophages were reported to have more than a hundred-fold increase in ISG15 mRNA levels, with the measured intracellular protein concentration reaching >700 ng/mL (∼0.5 µM in nucleus) [19]. Similarly, mRNA transcript levels of the E1 and E3 enzymes, Ube1L and HERC5, increased to 4 and 40 times the basal level during SARS-CoV-2 infection, respectively. An 8-fold increase for UbcH8 transcripts was also observed under the same conditions [19]. Data regarding molar concentration of UbcH8 is not available, however single cell RNA data for macrophages reports 64.8 nTPM (normalized transcript per million) and 138.4 nTPM, for UbcH8 and ISG15, respectively [20, 21, 22, 23]. Hence, we consider that intracellular UbcH8 concentration might be close to that of ISG15.

Considering that the CSP data indicates a possible conformational change in the E1 and E3 binding sites on UbcH8 associated with dimerization, it is also possible that elevated levels of E1 and E3 within the cell promote the dimeric state and thus increase E2 activity. Our results demonstrate that UbcH8, the E2 enzyme specific for ISGylation, may also form dimers at near physiological concentrations due to effects of molecular crowding and excluded volume effect. Although our experiments were performed at high concentrations, we believe that UbcH8 dimerization can also occur at lower concentrations within cells due to the effects of crowding and excluded volume. The crowded cellular environment effectively increases the local concentration of UbcH8 molecules, bringing them into closer proximity and enhancing the likelihood of interactions that can lead to dimerization. Additionally, the excluded volume effect limits the available space for UbcH8 molecules, further promoting interactions and potentially favoring dimer formation over other conformations. Locally high concentrations of proteins can occur due to macromolecular crowding, which can make protein-protein complex formation reactions more thermodynamically favorable [24]. This effect can modulate the behavior of proteins in the intracellular environment, where numerous macromolecules are present, and provide suitable conditions for oligomer formation at low molar concentrations.

This suggests that the ISGylation process may also involve dimerization for regulating the complicated interactions of E1, E2 and E3 enzymes. This study highlights the importance of understanding the oligomeric state and behavior of proteins in solution to gain insights into their biological function and regulation. Moreover, our work emphasizes the usefulness of SAXS and NMR techniques in elucidating protein structures and interactions in solution, which can complement crystallographic studies and provide a more biologically relevant context.

Future studies may focus on exploring the functional implications of UbcH8 dimerization in the ISGylation process, such as the effects on substrate specificity, E1 and E3 enzyme interactions, and the kinetics of ISGylation. Additionally, the molecular mechanisms underlying the observed concentration-dependent behavior of UbcH8 and the role of post-translational modifications in modulating its oligomeric state could be investigated further. These studies will contribute to a more comprehensive understanding of the regulation and function of UbcH8 in the context of ISGylation and its broader implications in various diseases, including viral and bacterial infections, cancer, and autoimmune disorders.

## MATERIALS AND METHODS

### Protein expression and purification

Three alanine residues followed by the coding sequence of the UbcH8 protein are inserted in the 5’ BamH1/ 3’ EcoR1 restriction sites of the pGEX-4T3 plasmid. Three alanine residues are inserted between the GST and UbcH8 protein sequence in order to increase the binding efficiency and also to provide a better cleavage upon thrombin treatment during the purification of the protein sample. The final coded amino acid sequence was:MSPILGYWKIKGLVQPTRLLLEYLEEKYEEHLYERDEGDKWRNKKFELMGLEFPNL PYYIDGDVKLTQSMAIIRYIADKHNMLGGCPKERAEISMLEGAVLDIRYGVSRIAYSKDFETLKVDFLSKLPEMLKMFEDRLCHKTYLNGDHVTHPDFMLYDALDVVLYMDPMCLDAFPKLVCFKKRIEAIPQIDKYLKSSKYIAWPLQGWQAFGGGDHPPKSDLVPRGSAAAMASMRVVKELEDLQKKPPPYLRNLSSDDANVLVWHALLLPDQPPYHLKAFNLRISFPPEYPFKPPMIKFTTKIYHPNVDENGQICLPIISSENWKPCTKTCQVLEALNVLVNRPNIREPLRMDLADLLTQNPELFRKNAEEFTLRFGVDRPS* pGEX-4T3 GST-AAA-UbcH8 plasmid was transformed into Rosetta2 *E. Coli* expression cells, plated on LB-ampicillin-chloramphenicol, and grown overnight at 37 °C. The next morning, colonies were picked from the agar plate and inoculated into 10 ml LB-ampicillin-chloramphenicol medium. The culture was grown overnight at 37 °C at 110 rpm. The overnight culture was transferred into 1 L LB medium and incubated at 37°C. After OD_595_ exceeded 0.3, temperature was lowered to 18°C and protein production was induced at OD_595_ 0.8 by the addition of 0.4mM IPTG. Cells were harvested 18 hours after induction by centrifugation at 2000 RCF for 1 hour.

Harvested cells were resuspended in lysis buffer (500mM NaCl, 50mM Tris, 0.1% (v/v) Triton X-100, 5% (v/v) glycerol, 1mM DTT, pH=7.5), sonicated, and centrifuged at 20K RCF for 1 hour to remove insoluble debris. The obtained supernatant was loaded to a GST affinity column equilibrated with 20mM Tris (pH 7.5), 150mM NaCl, 1mM DTT. Non-specific proteins were washed with the same buffer and the protein was eluted with 30mM glutathione, 20mM Tris (pH 7.5), 150mM NaCl, 1mM DTT. For cleavage of the GST tag, 1:100 thrombin enzyme was added to the eluted protein and dialyzed in 20mM Tris (pH 7.5), 150mM NaCl, 1mM DTT solution overnight to eliminate excess glutathione. For separation of the GST tag, reverse GST chromatography was applied. Unbound free UbcH8 was collected and purified by size exclusion chromatography using 20mM Tris (pH 7.5), 150mM NaCl, 1mM DTT buffer. Unlabeled ISG15N13Y/C78S protein is also expressed and purified as described above for NMR titration studies.

### SAXS Data Collection

All SAXS data were collected at home source SAXSpoint 5.0 (Anton Paar GmbH) as described before [25]. Sample/detector distance (SDD) was 1600 mm for SAXS experiments. All measurements took place at 10 °C. Data was collected in one hour session(1-minutes long 6 frames) for each measurement. The scattering curves were checked for radiation damage and no damage was detected after the superimposition of each 10 minute data collection intervals..

### SAXS Data Processing and Modeling

At first, the scattering pattern of all samples were visually inspected in the Primus program of ATSAS 3.0 for any possible issues with the measurement [11]. The radius of gyration (Rg) was calculated using Guinier’s equation and inverse Fourier transform by Primus. Distance distribution function P(r) and the maximum particle diameter (Dmax) was calculated by GNOM [26]. After estimating the molecular weight of the model DAMMIF (ab initio) is used to generate 5 independent low resolution models from the data [27]. DAMAVER and DAMMIN then averaged, clustered, and optimized these 5 distinct solutions to form the final ab-initio shape [28]. SASpy plug-in for PyMOL was used to superimpose the homology modeled structure of the protein [29, 30].

### 15N Labeled Protein Expression and Purification

pGEX-4T3 GST-AAA-UbcH8 plasmid containing bacteria were grown overnight in LB medium at 37°C and transferred into 50 mL ^15^N labeled M9 media the next day. Following 4 hours of incubation at 37°C, cells were transferred into 1L M9 media. After OD_595_ exceeded 0.3, temperature was lowered to 18°C and protein production was induced at OD_595_ 0.8 by the addition of 0.4mM IPTG. The medium contained 33.7mM Na_2_HPO_4_, 22 mM KH_2_PO_4_, 8.55 mM NaCl, 9.35 mM ^15^N labeled NH_4_Cl, 1mM MgCl_2_, 0.3mM CaCl_2_, and 7 mg/L FeCl_2_-4H_2_O. Cells were harvested 18 hours after induction by centrifugation at 2000 RCF for 1 hour.

Harvested cells were resuspended in lysis buffer (500mM NaCl, 50mM Tris, 0.1% (v/v) Triton X-100, 5% (v/v) glycerol, 1mM DTT, pH=7.5), sonicated, and centrifuged at 20K RCF for 1 hour to remove insoluble debris. The obtained supernatant was loaded to a GST affinity column equilibrated with 38.39 mM Na_2_HPO_4_, 11.61 mM KH_2_PO_4_ (pH 7.4), 100 mM NaCl, 1mM DTT. Non-specific proteins were washed with the same buffer and the protein was eluted with 30mM glutathione, 38.39 mM Na_2_HPO_4_, 11.61 mM KH_2_PO_4_ (pH 7.4), 100 mM NaCl, 1mM DTT. For cleavage of the GST tag, 1:100 thrombin enzyme was added to the eluted protein and dialyzed in 38.39 mM Na_2_HPO_4_, 11.61 mM KH_2_PO_4_ (pH 7.4), 100 mM NaCl, 1mM DTT solution overnight to eliminate excess glutathione. For separation of the GST tag, reverse GST chromatography was applied. Unbound free UbcH8 was collected and purified by size exclusion chromatography using 38.39 mM Na_2_HPO_4_, 11.61 mM KH_2_PO_4_ (pH 7.4), 100 mM NaCl, 1mM DTT buffer.

### NMR Data Acquisition and analysis

UbcH8 was concentrated to 0.150 mM and 0.287 mM. 10% D2O (final concentration) containing 1mM DSS was added to obtain a final sample volume of 600 µL. For NMR titration; UbcH8 was concentrated to 0.300 mM and from a series of 1H, ^15^N-HSQC spectra acquired from samples with ^15^N-UbcH8 and unlabeled ISG15 molar ratios of 1:0.25, 1:0.5, 1:1, 1:2, 1:4. All NMR data acquisition process was completed using 500 MHz Bruker Ascend magnet equipped with Avance NEO console and BBO double resonance room temperature probe at Koç University n^2^STAR NMR Facility. 2-D ^1^H-^15^N HSQC spectra were recorded with 50% non-uniform sampling (NUS) at 298 K with a 1H spectral width of 14 ppm (1024 data points in t2) and a ^15^N spectral width of 32 ppm (64 data points in t1). The 2D data was processed by NMRPipe [31] and analyzed using NMRFAM-SPARKY [32]. The combined ^1^H-^15^N chemical shift perturbations were calculated using equation ΔδAV =[(Δδ 1H * 5)^2^ + (Δδ 15N)^2^]^1/2^.

1D TRACT experiments [11] were collected with 1024 complex points and 1.5 s recycle delay. Relaxation rates for ^15^N TROSY and anti-TROSY components were determined from spectra intensity values integrated over 9.2 to 8.6 ppm at eight relaxation delays: 30, 60, 90, 120, 150, 180, 210, 240, 270 and 300 ms. Each relaxation rate and its uncertainty was estimated by fitting the integrated values and time delays to a single parameter exponential decay model using Bayesian Parameter Estimation. Each TROSY and anti-TROSY relaxation rate was then used to estimate rotational correlation time (*π_c_*) using the algebraic method [12] where we assumed an order parameter (*O*^2^) of 0.8.

## Author Contributions

**Çağdaş Dağ:** Conceptualization, Methodology, Supervision, Funding acquisition, Writing-Reviewing and Editing, Writing-Original draft preparation **Joshua J. Ziarek:** Supervision, Conceptualization, Writing-Reviewing and Editing, Writing-Original draft preparation **Arthur L. Haas:** Conceptualization, Supervision **Volker Dötsch:** Supervision, Writing-Reviewing and Editing **Emine Sonay Elgin:** Conceptualization, Supervision, Writing-Reviewing and Editing **Kerem Kahraman**: Investigation, Formal analysis, Visualization. **Scott A. Robson:** Investigation, Formal analysis, Visualization, Writing-Reviewing and Editing **Oktay Göcenler:** Formal analysis, Investigation Writing - Original Draft, Visualization **Cansu D. Tozkoparan:**Visualization, Formal analysis **Jennifer M. Klein:** Investigation, **Cansu M. Yenici:** Investigation

## Protein Accession IDs

**Q96LR5:** UbcH8 / UB2E2_HUMAN / Ubiquitin-conjugating enzyme E2 E2

**P08515:** Glutathione S-transferase class-mu 26 kDa isozyme / GST

## ACKNOWLEDGEMENT

CD acknowledges support from TÜBİTAK (Project No: 120Z594, 122Z747). JJZ acknowledges support from National Institutes of Health grant R35GM143054. The authors acknowledge the use of the services and facilities of n^2^STAR-Koç University Nanofabrication and Nanocharacterization Center for Scientific and Technological Advanced Research.

