## Supplementary material for "Characterizing the monomer-dimer equilibrium of UbcH8/Ube2L6: A combined SAXS and NMR study": Supp Figs

### TOC

**Table S1……………………………………..S3**

**Table S2……………………………………..S4**

**Figure S1……………………………………..S5**

**Figure S2……………………………………..S6**

**Figure S3……………………………………..S7**

**Figure S4……………………………………..S8**

**Figure S5……………………………………..S9**

**Figure S6……………………………………..S10**

**Figure S7……………………………………..S11**

**Figure S8……………………………………..S12**

**Table S1.** Molecular Size Parameters of GST-UbcH8 and UbcH8 dimer Obtained from SAXS Data Analysis.

| Sample | $R_g$ (nm)[a] | Dmax (nm)[b] | Vp (Å$^3$)[c] | MW (kDa)[d] | Vc (kDa)[e] | Qp (kDa)[f] |
|---|---|---|---|---|---|---|
| GST-UbcH8 | 3.37 | 8 | 8919 | 41.604 | 41.737 | 42.842 |
| UbcH8 dimer | 4.402 | 6.2 | 5835 | 39.547 | 38.766 | 44.169 |

[a]Radius of gyration, [b]Maximum Dimension, [c]Porod Volume, [d]Molecular Weight, [e]Volume of Correlation, [f]Porod invariant

**Table S2.** Fit quality for superimposition of E2 crystal structures onto UbcH8 dummy atom model. The UbcH8 dummy atom model was calculated from the SAXS data using DAMMIF. The fineness and normalized spatial discrepancy (NSD) of the superimposition obtained using the SASpy Pymol extension.

| | **3WE5** | **6QH3** | **6QHK** | **6S98** |
|---|---|---|---|---|
| **Fineness** | 1.549 | 1.370 | 1.030 | 1.043 |
| **Normalized Spatial Discrepancy (NSD)** | 2.2294 | 3.6498 | 4.9168 | 5.3306 |

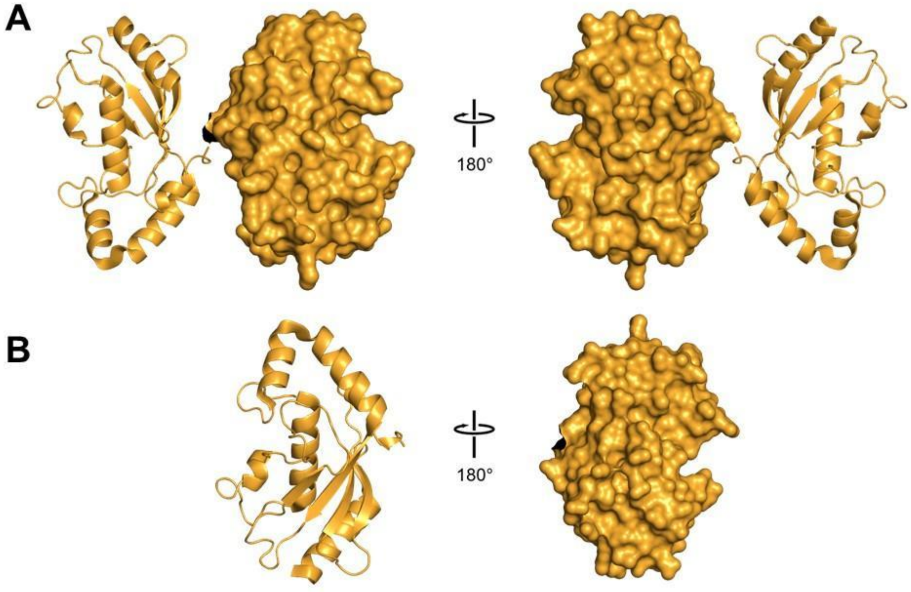


**Figure S1. Previously-determined UbcH8 crystal structures.** Cartoon and surface representation of UbcH8 protein in the (A) dimeric form (PDB ID:1WZV) and (B) monomeric form (PDB ID:1WZW)

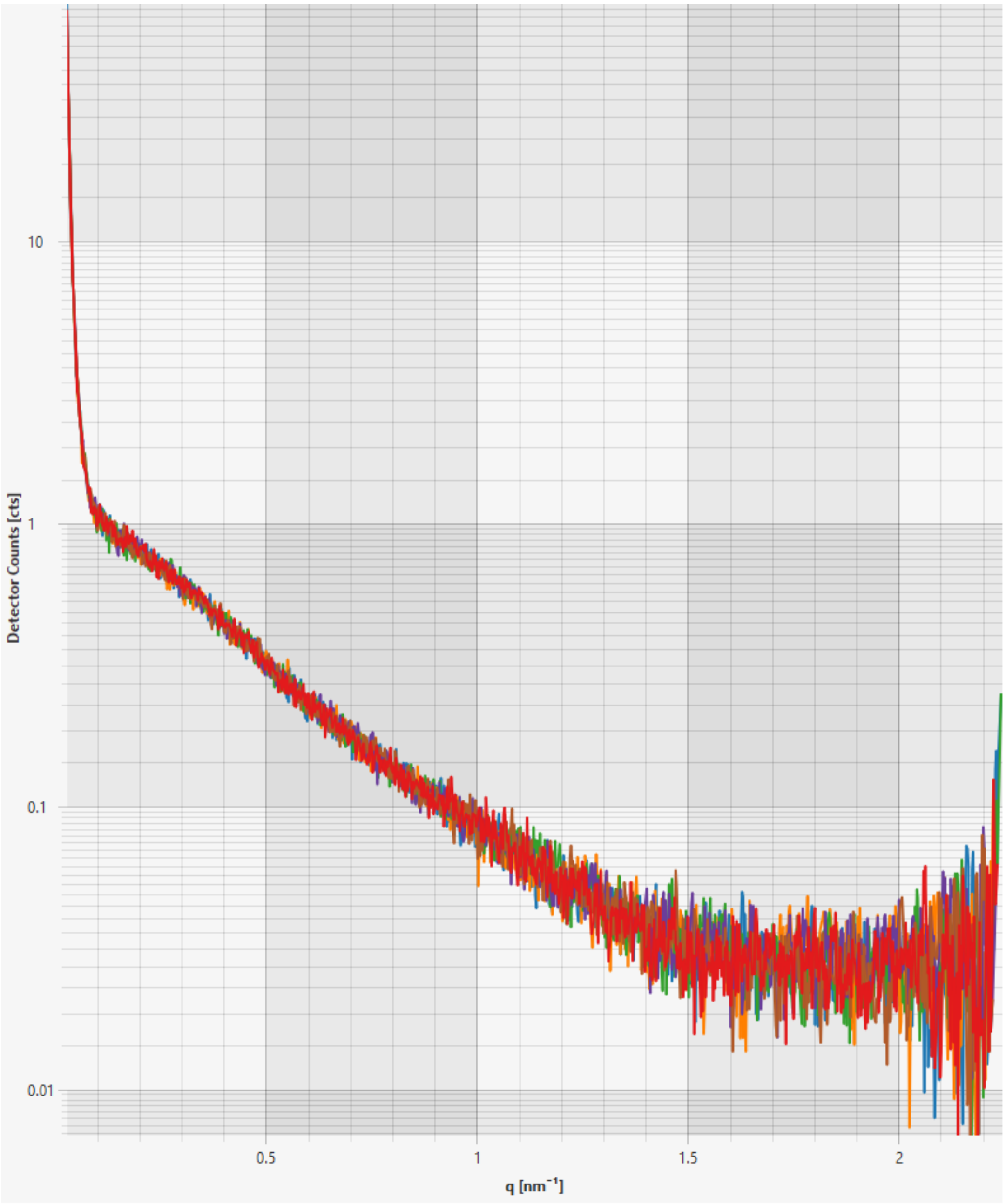


**Figure S2. Superimposition of six, 10 minute SAXS frames.** The data indicates that little to no radiation damage occurred during the total acquisition period.

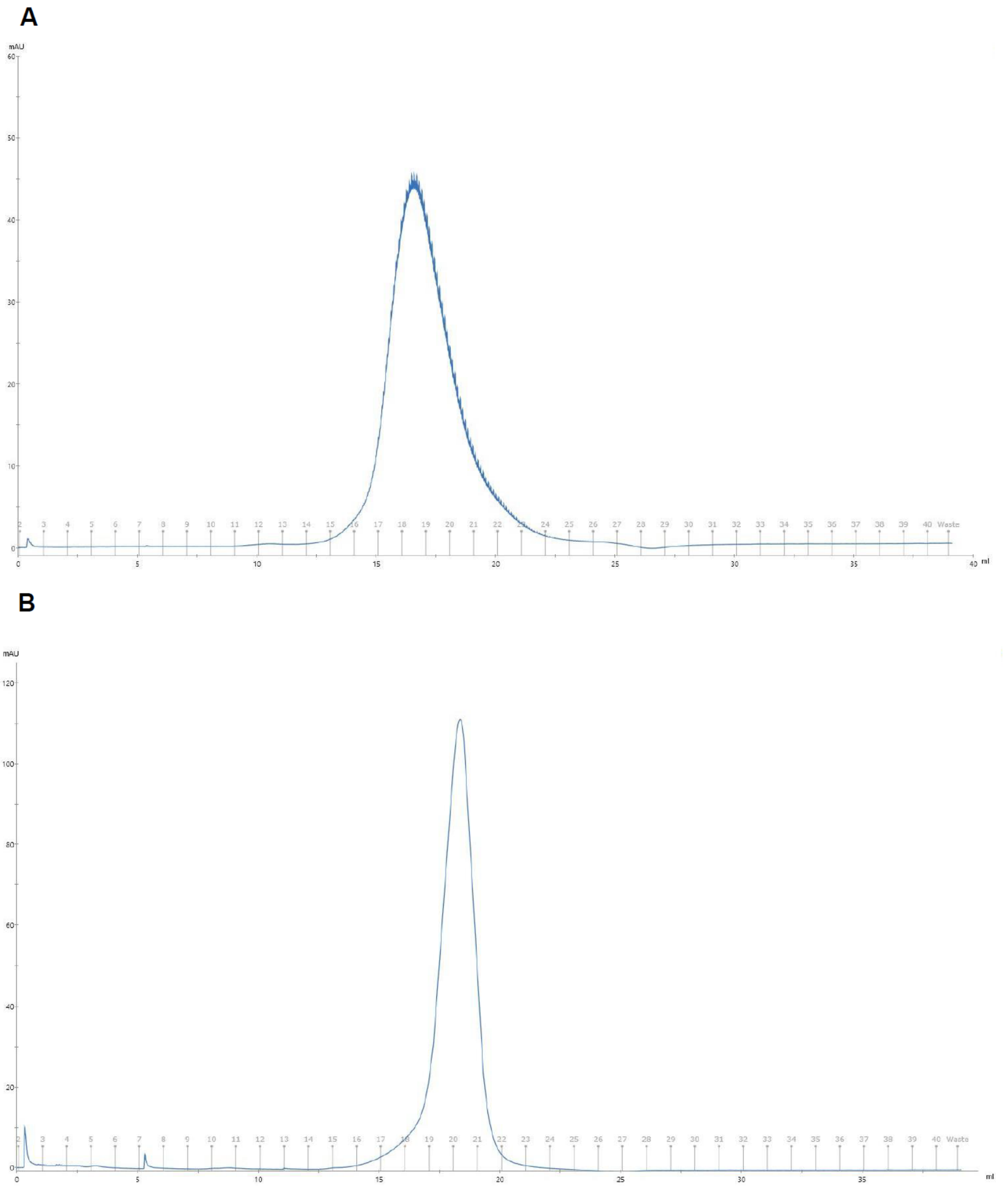


**Figure S3.** SEC chromatograms of (A) UbcH8 (B) 12.2 kDa monomeric protein. S200 16/600 SEC column was used with identical run parameters.

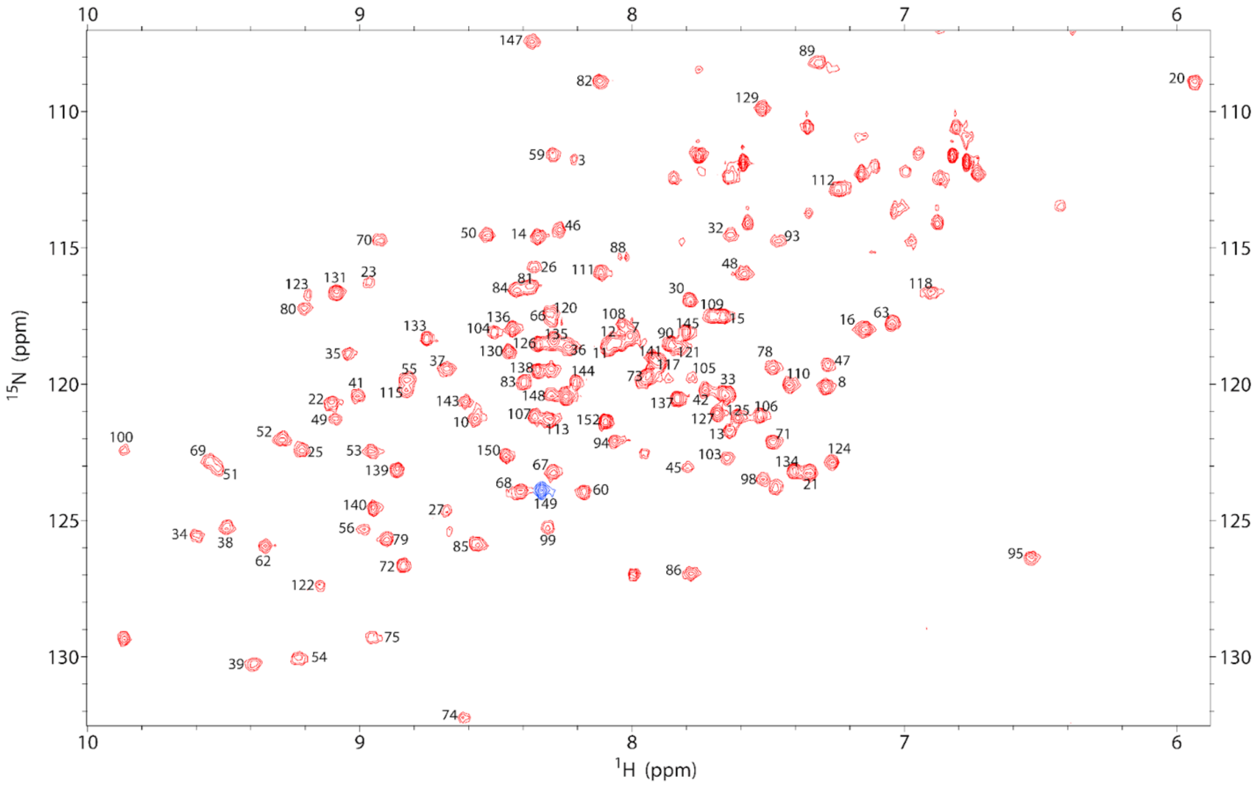


**Figure S4. $^{1}$H-$^{15}$N HSQC spectrum of UbcH8 collected at 25°C.** Residue numbers are indicated on each peak. Residue 149 (blue) is located at the dimerization interface of the crystal structure. Resonances were assigned by visual inspection using BMRB 16321. Data were collected at 300 μM on a 11.7 T Bruker NMR spectrometer.

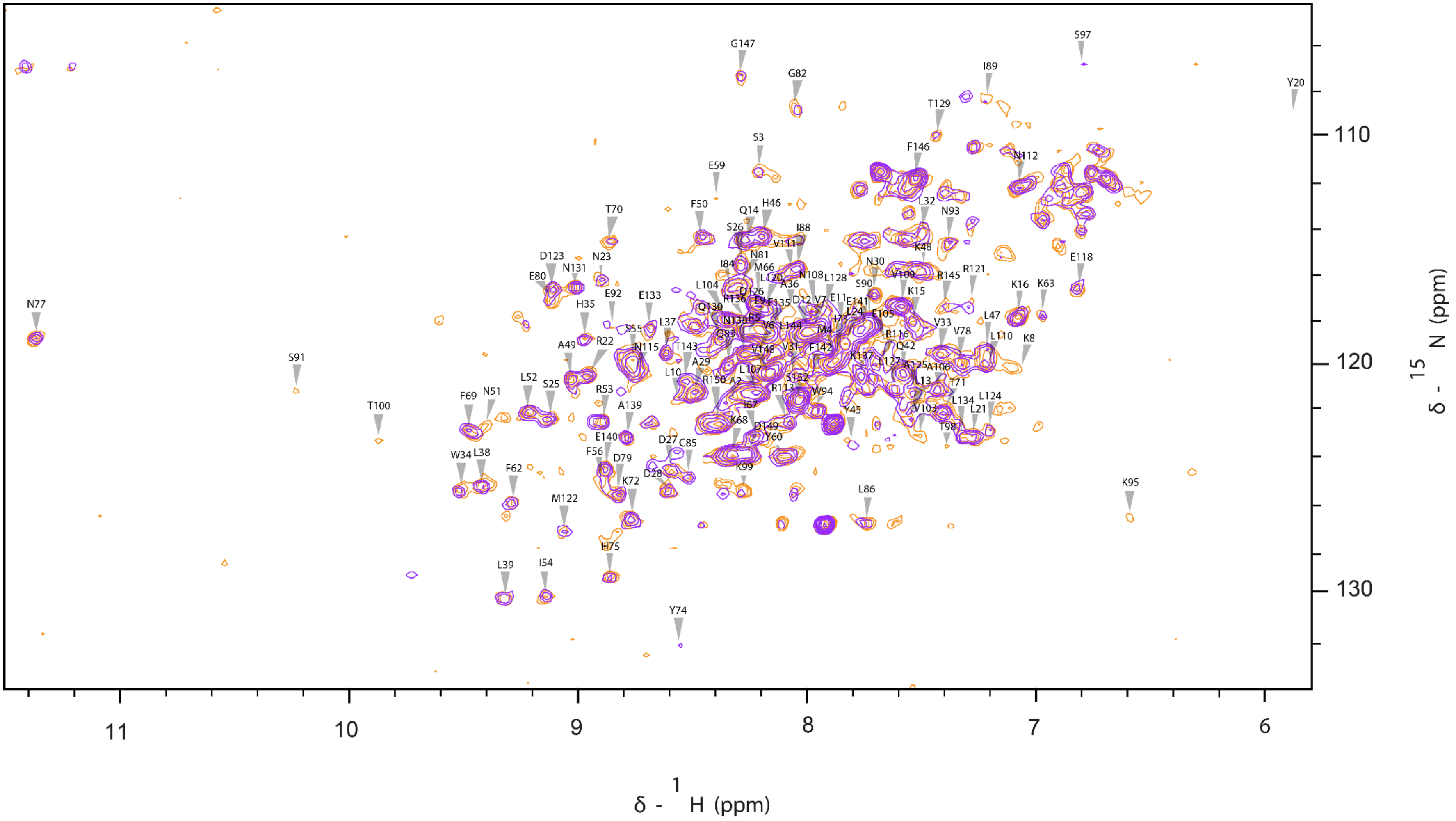


**Figure S5. $^{1}H$-$^{15}N$ HSQC superimposed spectra of UbcH8 collected at 25°C.** Orange and purple indicate data collected at 150µM and 300µM, respectively.

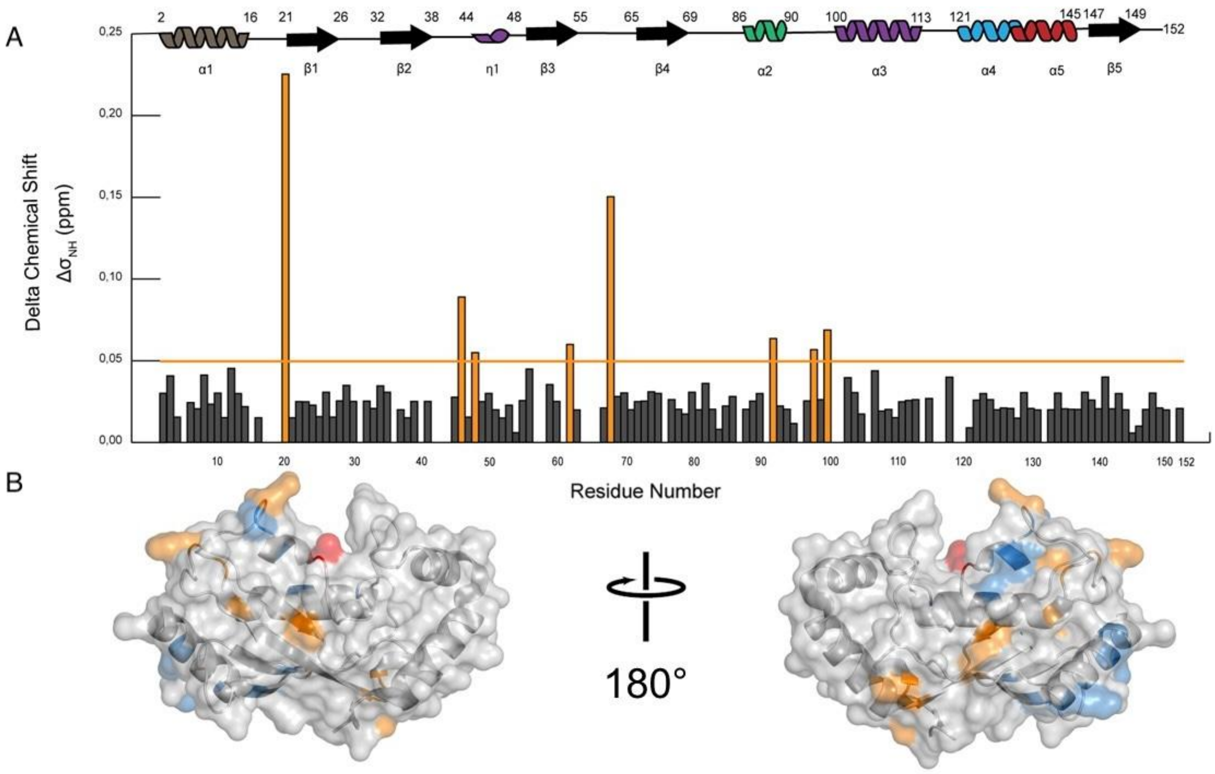


**Figure S6. $^{1}H$-$^{15}N$ Chemical shift perturbations of $^{15}N$ labeled UbcH8 titrated with ISG15.** A) The combined $^{1}H$-$^{15}N$ chemical shift perturbations were calculated for each residue. The residues colored orange possessed chemical shift perturbations larger than the threshold (orange line). Residues with no bars were not observed at any concentration. B) Residues with chemical shift perturbations larger than the threshold were mapped onto the UbcH8 monomeric crystal structure (PDB 1WZW) labeled with orange. Peaks with significant intensity changes are indicated in blue, while the catalytic cysteine is labeled red.

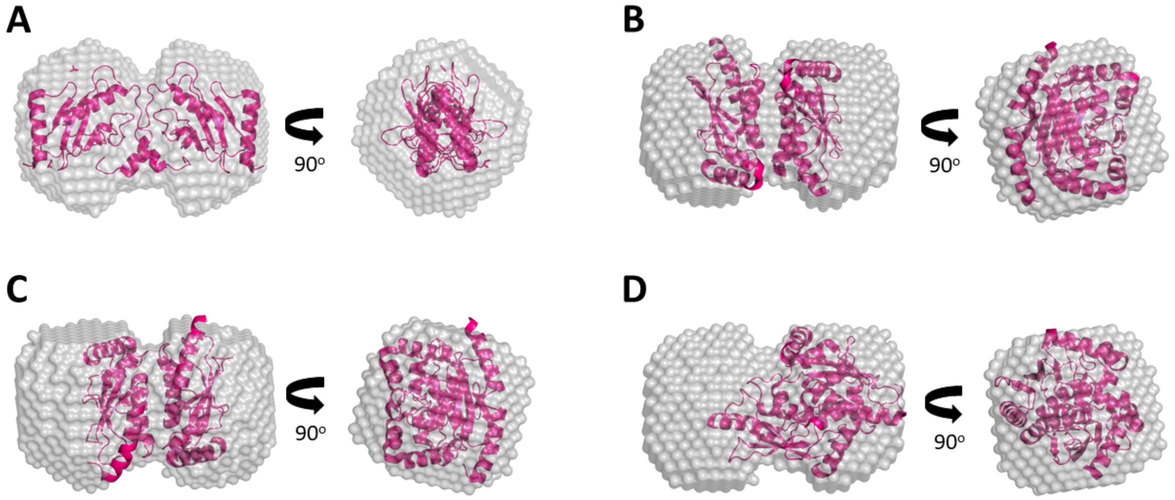


**Figure S7. Comparison of free UbcH8 dummy atom model with previously-determined dimeric E2 enzyme crystal structures.** (A) Ubiquitin conjugating enzyme E2 UbcA1 from Agrocybe aegerita (PDB: 3WE5). (B) The catalytic domain of the human ubiquitin-conjugating enzyme UBE2S C118M (PDB: 6QH3). (C) The catalytic domain of the human ubiquitin-conjugating enzyme UBE2S (PDB: 6QHK). (D) The catalytic domain of wild-type UBE2S (PDB: 6S98). The free UbcH8 dummy atom model was calculated by DAMMIF.

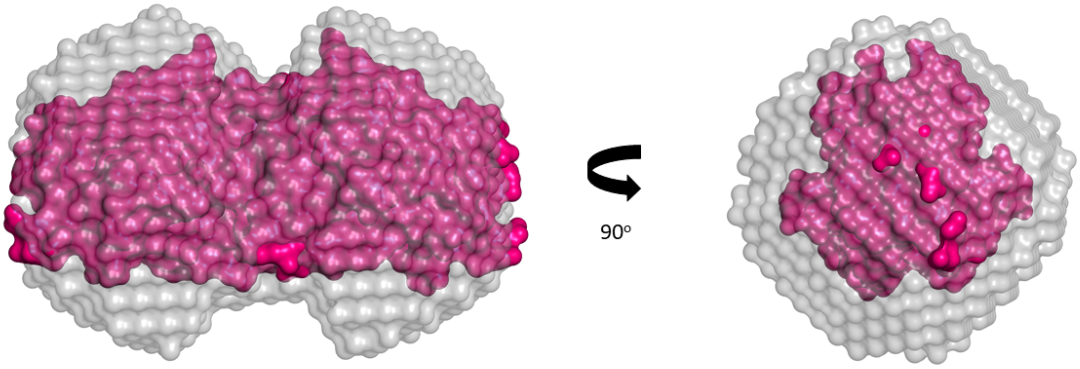


**Figure S8. UbcA1 dimer crystal structure (pdb:3WE5) fitted into the DAMMIF dummy atom model** Crystal structure of ubiquitin conjugating enzyme E2 UbcA1 from Agrocybe aegerita (PDB: 3WE5) shown as surface (pink) placed into the dummy model of UbcH8 dimer (Free UbcH8, gray) obtained after the DAMMIF analysis of the SAXS data, using Pymol plugin SasPY.
